# The presynaptic vesicle cluster transitions from a compact to loose organization during long-term potentiation

**DOI:** 10.1101/2024.11.01.621529

**Authors:** Guadalupe C. Garcia, Thomas M. Bartol, Lyndsey M. Kirk, Priyal Badala, Kristen M. Harris, Terrence J. Sejnowski

**Author notes:** Co-communicating authors (Kristen M. Harris, Terrence J. Sejnowski).

## Abstract

Functional and structural elements of synaptic plasticity are tightly coupled, as has been extensively shown for dendritic spines. Here, we interrogated structural features of presynaptic terminals in 3DEM reconstructions from CA1 hippocampal axons that had undergone control stimulation or theta-burst stimulation (TBS) to produce long-term potentiation (LTP). We reveal that after LTP induction, the synaptic vesicle (SV) cluster is less dense, and SVs are more dispersed. The distances between neighboring SVs are greater in less dense terminals and have more SV-associated volume. We characterized the changes to the SV cluster by measuring distances between neighboring SVs, distances to the active zone, and the dispersion of the SV cluster. Furthermore, we compared the distribution of SVs with randomized ones and provided evidence that SVs gained mobility after LTP induction. With a computational model, we can predict the increment of the diffusion coefficient of the SVs in the cluster. Moreover, using a machine learning approach, we identify presynaptic terminals that were potentiated after LTP induction. Lastly, we show that the local SV density is a volume-independent property under strong regulation. Altogether, these results provide evidence that the SV cluster is undergoing a transition during LTP.

## Introduction

Synaptic plasticity is the process by which synapses are modified in structure and function in response to activity, thereby generating a change in synaptic efficacy. Thus, synaptic plasticity alters the strength of neural circuits and is fundamental for animals to learn and adapt to environmental changes. Hence, understanding the interplay of structural and functional changes during learning may provide new insights into how the strength of synapses is regulated.

Long-term potentiation (LTP) is a cellular model of learning that produces a persistent strengthening of synapses lasting hours to days [1]. Converging evidence from recent studies reveals novel presynaptic mechanisms contributing to LTP [2]. These include changes in axonal bouton structure following high-frequency stimulation [3]. Moreover, at CA1 terminals several structural changes occur following 2 hours of theta-burst stimulation (TBS) to produce LTP. It has been shown that the number of nondocked SVs decreases [4, 5] while the boutons’ surface area substantially increases [6] 2 hours after the induction of LTP. The reduction of nondocked SVs is considerable in presynaptic terminals that contain mitochondria [7]. Furthermore, following 2 h of TBS there is an increase in mitochondrial localization to synapses and volume of these organelles [7].

Redistribution of SVs after LTP was also observed in electron microscopy (EM) images [8]. A migration of SV towards the active zone (AZ) was suggested, accompanied by a decrease of the SV density following repetitive hippocampal stimulation [8]. Depletion of SVs was also reported in the entorhinal cortex (EC) 10-60 min after 30 seconds of tetanic stimulation [9].

An additional level of synaptic protein organization has been recently identified: the formation of biological condensates [10]. In presynaptic terminals, the AZ and SV cluster components have been shown to form condensates [11]. In particular, condensates of synapsin I have been implicated in clustering lipid vesicles in vitro [12] and SVs in vivo [13, 14]. SVs are known to interact with several phosphoproteins [15, 16, 17] that regulate the state of the SV cluster. The phosphorylation of some of these proteins by synaptic activation has been suggested to control the clustering of SVs [17, 15].

In this work, we employed quantitative methods and a simple mathematical model to answer two questions: Is the structure of the SV cluster altered after induction of LTP in hippocampal boutons synapsing with CA1 dendritic spines? Second, what are the functional implications of any changes in the structure or composition of the SV clusters? To answer these questions, we developed methods to characterize the structure of the SV cluster. The outcomes reveal that the structure of the SV cluster is altered after LTP induction: the SV cluster is less dense, more dispersed, and more similar to a random distribution after LTP induction, while other properties remain unaltered as the position of the SV cluster center and the distance from the SVs to the AZ. Moreover, we show that the local SV density is a volume-independent property that changes after LTP induction. Finally, we combined morphological information with a simple mathematical model to predict the impact of the change in SV density. We provide evidence supporting the hypothesis that SVs in the cluster move faster after the induction of LTP. Altogether, these results provide evidence that the SV cluster is undergoing a transition during LTP.

## Results

### LTP augments bouton and mitochondrial volume relative to control conditions

Several structural changes occur at CA1 presynaptic terminals following TBS to produce LTP [4, 5, 18]. In particular, it has been shown that the number of nondocked SVs decrease [7, 6], while the surface area of the boutons increases [6] and presynaptic mitochondria enlarge [7] following 2h of TBS.

Here, we examined 3D reconstructions of axons in adult Long-Evans rats stimulated under control conditions and after 2h of TBS to produce LTP (Fig. 1a-d). Detailed 3D reconstructions of axons and mitochondria were generated from curated 2D traced contours (Fig. 1a-d). The position of nondocked SVs was also assessed, and spherical objects were created at their locations.

**Figure 1.**
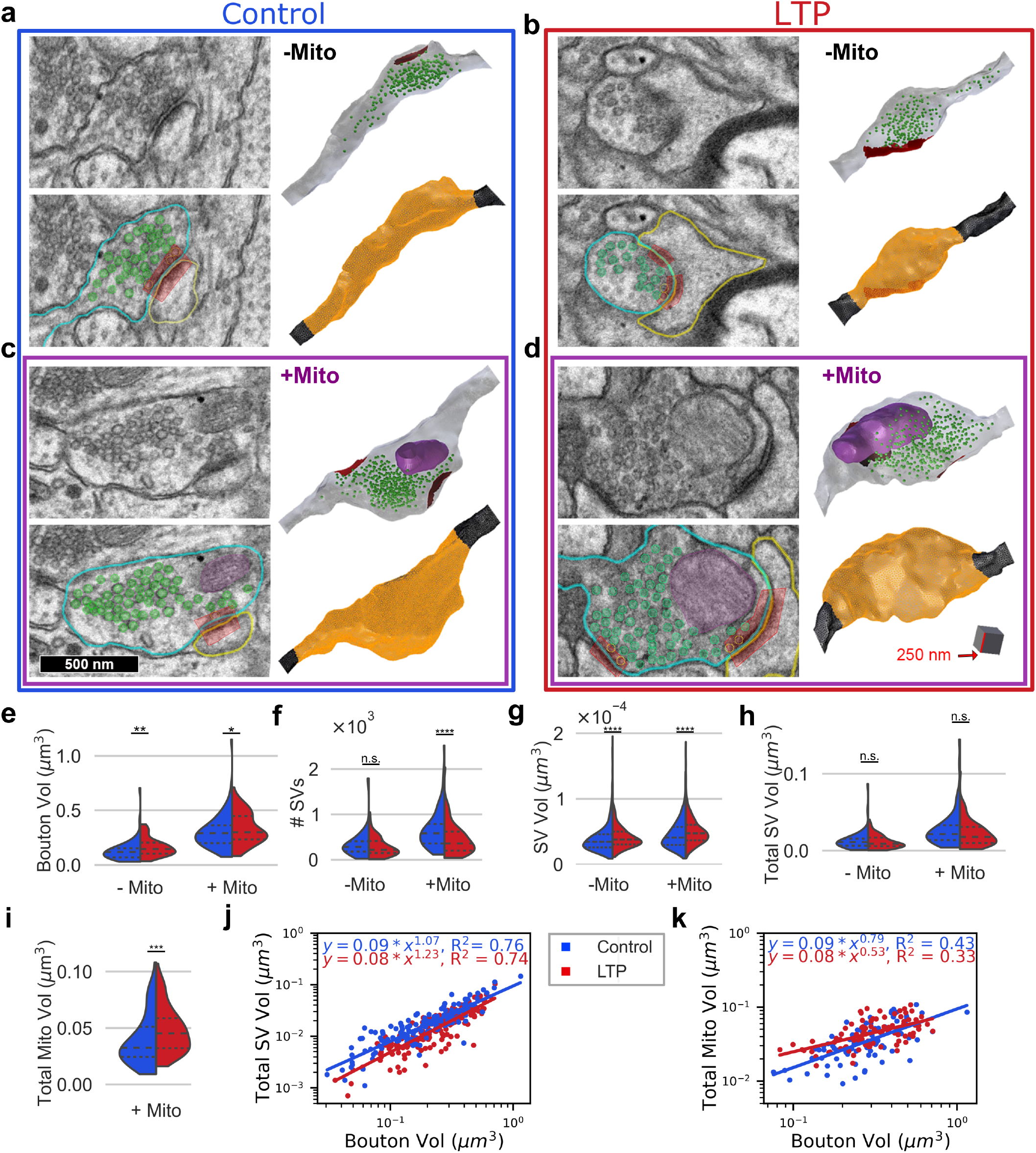
LTP augments bouton and mitochondrial volume relative to control conditions. (a-d) EM images of representative boutons and their 3D reconstructions: (a-c) Under control conditions in blue; (b-d) under LTP in red. (a-b) boutons without mitochondria (-Mito) and (c-d) boutons with mitochondria (+Mito). Here, SVs are represented in green, mitochondria in magenta, and active zones in red. The surface area of the bouton is highlighted in orange in the 3D reconstructions. (e) The bouton volume increases after LTP induction. (f) The number of SVs per bouton decreases after LTP. (g) However, the individual SV volume sampled from boutons increases after LTP induction. Because of the net effect of (f) and (g), (h) the total SV volume remains unaltered after LTP. (i) Total mitochondrial volume increases after LTP. (j-k) Correlation between the bouton volume with (j) total SV volume and (k) total mitochondrial volume. Violin plots (e-i) show quartiles 25th, 50th, and 75th. Significant differences are indicated as ^*^p < 0.05, ^**^p < 0.01, ^***^p < 0.001, and ^****^p < 0.0001. Medians, standard deviations, and p-values are in Table 1.

We precisely quantified the bouton volumes in 3D reconstructions of axons. After 2h of LTP induction boutons enlarge (Fig. 1e). We separately analyzed boutons lacking mitochondria (-Mito) and with mitochondria (+Mito) since differences were previously reported [7]. Following TBS, boutons lacking mitochondria are larger on average than those under control conditions (Fig. 1e, p < 0.0019). A similar result is observed for boutons with mitochondria (Fig. 1e, p < 0.04). Moreover, boutons containing mitochondria are significantly larger than those lacking organelles under both conditions (under control conditions, p < 2.62 × 10^−14^; and under LTP conditions, p < 4.40 × 10^−16^). Medians and standard deviations of the computed volumes are in Table 1.

**Table 1:** Hippocampal CA1 boutons, mitochondria, and SVs dimensions. Medians and standard deviations are shown. (-M) boutons lacking mitochondria and (+M) boutons with mitochondria.

| Feature | Control (-M) | LTP (-M) | p-val | Control (+M) | LTP (+M) | p-val |
| --- | --- | --- | --- | --- | --- | --- |
| Number of boutons | 61 | 67 |  | 81 | 88 |  |
| Bouton Volume ($\mu\text{m}^3$) | $0.1 \pm 0.1$ | $0.15 \pm 0.08$ | 0.0019 | $0.29 \pm 0.16$ | $0.30 \pm 0.14$ | 0.044 |
| Number of SVs | $285 \pm 312$ | $218 \pm 221$ | 0.12 | $578 \pm 434$ | $357 \pm 342$ | $4 \times 10^{-5}$ |
| SV Volume ($\mu\text{m}^3$) | $(3.5 \pm 0.2) \times 10^{-5}$ | $(3.8 \pm 0.2) \times 10^{-5}$ | $3 \times 10^{-10}$ | $(4.1 \pm 0.2) \times 10^{-5}$ | $(4.7 \pm 0.2) \times 10^{-5}$ | $3 \times 10^{-7}$ |
| Total SV Vol ($\mu\text{m}^3$) | $0.01 \pm 0.01$ | $0.009 \pm 0.009$ | 0.32 | $0.02 \pm 0.02$ | $0.018 \pm 0.018$ | 0.05 |
| Total Mito Vol ($\mu\text{m}^3$) | 0 | 0 | 0 | $0.03 \pm 0.02$ | $0.05 \pm 0.02$ | 0.0006 |
| Dis AZ ($\mu\text{m}$) | $0.20 \pm 0.08$ | $0.20 \pm 0.07$ | 0.4 | $0.25 \pm 0.07$ | $0.23 \pm 0.08$ | 0.15 |
| Dis Neigh SVs ($\mu\text{m}$) | $0.03 \pm 0.02$ | $0.043 \pm 0.018$ | $5 \times 10^{-16}$ | $0.026 \pm 0.006$ | $0.04 \pm 0.02$ | $7 \times 10^{-22}$ |
| Dis SV CC ($\mu\text{m}$) | $0.15 \pm 0.05$ | $0.19 \pm 0.06$ | $5 \times 10^{-7}$ | $0.21 \pm 0.06$ | $0.14 \pm 0.01$ | $4.8 \times 10^{-3}$ |
| $\sigma_{xyz}$ ($\mu\text{m}$) | $0.19 \pm 0.06$ | $0.25 \pm 0.07$ | $1 \times 10^{-8}$ | $0.26 \pm 0.07$ | $0.3 \pm 0.1$ | $8 \times 10^{-5}$ |
| Clu-associated Vol ($\mu\text{m}^3$) | $0.05 \pm 0.08$ | $0.08 \pm 0.07$ | $8 \times 10^{-5}$ | $0.12 \pm 0.01$ | $0.14 \pm 0.01$ | $5 \times 10^{-2}$ |
| SV-associated Vol ($\mu\text{m}^3$) | $(8 \pm 2) \times 10^{-5}$ | $(17 \pm 6) \times 10^{-5}$ | $1.3 \times 10^{-19}$ | $(8 \pm 2) \times 10^{-5}$ | $(16 \pm 8) \times 10^{-5}$ | $8 \times 10^{-5}$ |
| Glob SV Den ($\mu\text{m}^{-3}$) | $6249 \pm 2200$ | $3028 \pm 1087$ | $3 \times 10^{-16}$ | $4859 \pm 1344$ | $2616 \pm 908$ | $9 \times 10^{-21}$ |
| Loc SV Den ($\mu\text{m}^{-3}$) | $12259 \pm 3535$ | $5779 \pm 1849$ | $8 \times 10^{-20}$ | $11883 \pm 2423$ | $5779 \pm 1849$ | $6 \times 10^{-24}$ |

The number of SVs per bouton decreases after LTP (Fig. 1f), as was already reported [4]. However, the individual SV volumes sampled from boutons increase after LTP induction (Fig. 1g). Therefore, because of the net effect of (f) and (g) the total SV volume at each bouton (Fig. 1h) remains unaltered after LTP induction. On the contrary, the total mitochondrial volume increases after LTP induction (Fig. 1i) as was previously reported [7].

Next, we investigated how the total SV volume relates to the bouton volume (Fig. 1j). These quantities are highly correlated under control and LTP conditions. Still, we observe a lower total SV volume for a fixed bouton volume, which means boutons of the same volume under LTP have less total SV volume.

The total mitochondrial volume is moderately correlated with the bouton volume, and small differences are observed between control and LTP conditions (Fig. 1k), mainly due to the increment of mitochondrial volume for smaller boutons.

### SVs are spread out after LTP

To analyze the 3D spatial distribution of nondocked SV in boutons, we computed distances between SVs. First, we compute the distance from the SVs to the AZ (Fig. 2a). The median distance from the SVs to the AZ remains unaltered after LTP induction. Similar results are observed for single synaptic boutons and multisynaptic boutons (Fig. 1 in the supplementary material). The percentage of SVs closer to the AZ (with distance smaller than 150 nm) is also similar under control and LTP conditions (≈ 25% of the SVs are closer to the AZs). The median distance from the SVs to the AZ is slightly smaller for boutons without mitochondria (p < 0.001 under control conditions, and p < 0.007 under LTP).

**Figure 2.**
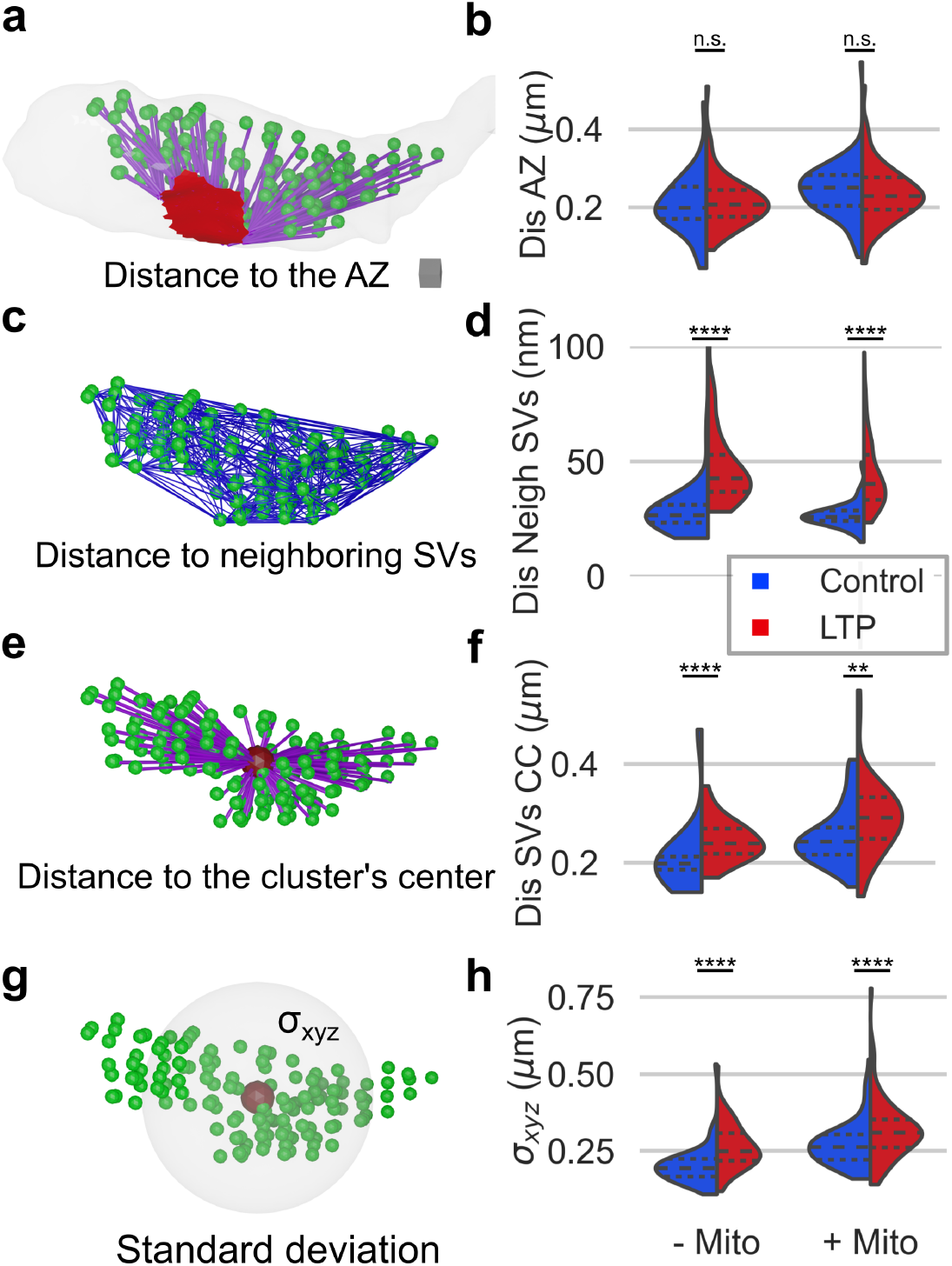
SVs are spread out after 2 hours of LTP induction. (a) Representative example of a bouton and the SVs (in green) after LTP induction. Distances from the SVs to the active zone are shown in purple, and the active zone in red. Scale cube = 0.001 µm^3^. (b) No differences in the distances from the SVs to the AZ are observed under LTP, on the left for boutons without mitochondria (-Mito) and on the right for boutons with mitochondria (+ Mito). (c) Delaunay triangulation for a representative SV cluster (in blue). We computed the mean distances to neighboring SVs (Dis Neigh SVs) at each bouton, considering neighboring SVs those connected in the Delaunay triangulation. (d) The mean distance between neighboring SVs increases under LTP, and the increment is similar for boutons with and without mitochondria. (e) Visual representation of the center of the SV cluster (CC, in dark red) and the distances from the SVs to the cluster’s center in purple (Dis SVs CC, in purple). (f) SVs are further away from the center of the SV cluster under LTP. (g) Visual representation of the standard deviation (σ_*xyz*_) of the position of the SVs from the cluster’s center –a measure of the SVs’ dispersion from the cluster’s center. (h) SVs are more spread out after LTP induction. Violin plots show quartiles 25th, 50th, and 75th. Significant differences are indicated as ^*^p < 0.05, ^**^p < 0.01, ^***^p < 0.001, and ^****^p < 0.0001. Medians, standard deviations, and p-values are in Table 1.

Next, we measured the pairwise distance between all SVs in the cluster and computed for each SV the mean distance to its neighbors. The neighbors were determined using the Delaunay triangulation for the SV cluster (Fig. 2c); connected SVs were considered neighbors. The distance between neighboring SVs is larger following LTP (Fig. 2d and Table 1). No significant differences between the distances were observed between boutons with and without mitochondria or between single synaptic boutons (SSB) and multi-synaptic boutons (MSB) (Fig 2 in the supplementary material).

We determined the position of the SV cluster’s center (as the mean of the XYZ coordinates of the SVs, Fig. 2e), and computed the distances from the SVs to the cluster’s center (Fig. 2e, in purple). The distances from the cluster’s center to the SVs increase after LTP induction (Fig. 2f).

To understand in more detail the spatial organization of SVs, we measured the dispersion of the SVs from its center (Fig. 2g). SVs are more dispersed from the cluster’s center after LTP induction (Fig. 2h). Moreover, the dispersion is larger for boutons with mitochondria (under control conditions p < 6.00 ×10^−11^ and under LTP p < 9.00 ×10^−5^).

Other measured quantities remained unaltered after LTP induction: the position of the SV cluster’s center relative to the AZ or the minimal distance from the SVs to mitochondria (Table 1).

### The volume of the SV cluster is maintained under LTP, and the SV density is reduced

Next, we asked if the volume of the SV cluster collapses due to the drop in SV number after LTP induction. For this, we measured the SV cluster volume, which we defined as the convex hull of the SVs (see methods and Fig. 3a). We corrected the volume by the estimated radius of the SVs and intersected it with mitochondria and the axons to make the estimation more precise (see methods). The cluster-associated volume (Clu-associated Vol) is the remaining volume after subtracting the SV volume from the minimal volume of the SV cluster (Fig. 3b). The cluster-associated volume increases for boutons without mitochondria after LTP induction (p-val < 8 × 10^−5^) and is maintained for boutons with mitochondria (Fig. 3c, p-val < 5 × 10^−2^).

**Figure 3.**
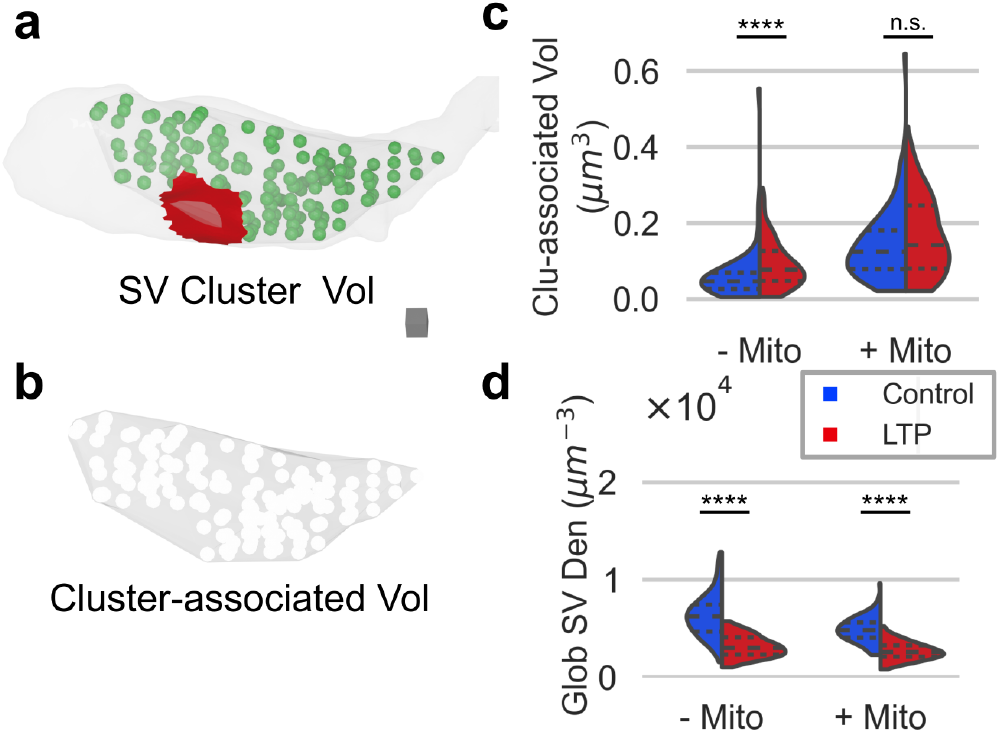
The volume associated with the SV cluster is maintained after LTP induction rather than collapsing with the loss of SVs. (a) Minimal volume of the SV cluster in 3D (the volume was calculated using the convex hull, see methods). Scale cube = 0.001 µm^3^. (b) The cluster-associated volume (Clu-associated Vol) is the remaining volume after subtracting the SV volume from the minimal volume of the SV cluster. (c) The cluster-associated volume increases in boutons without mitochondria (-Mito) and is maintained in boutons with mitochondria (+Mito) at 2 hours after LTP induction. Differences in the cluster-associated volume are observed for boutons without mitochondria (-Mito) and with mitochondria (+ Mito) under control conditions (in blue, p-val < 1.30 × 10^− 12^) and under LTP (in red, p-val < 1.50 × 10^− 6^). (d) The global SV density (Glob SV Den) decreases under LTP (the total number of SVs in the bouton divided by the cluster-associated volume). Violin plots show quartiles 25th, 50th, and 75th. Significant differences are indicated as ^*^p < 0.05, ^**^p < 0.01, ^***^p < 0.001, and ^****^p < 0.0001. Medians, standard deviations, and p-values are in Table 1.

We also calculated the global SV density (Glob SV Den), i.e. the number of nondocked SVs divided by the cluster-associated volume (Fig. 3d). The global SV density decreases for boutons without and with mitochondria (p-val < 3 ×10^−16^ and p-val < 9 ×10^−21^ respectively). Since many SVs are lost after LTP induction, and the volume of the SV cluster is maintained, it is not surprising to find a reduction in the SV density at the boutons. Altogether, these results show that boutons are densely packed under control conditions, and in particular, boutons without mitochondria are more densely packed.

We generated a partition of the SV cluster and associated a specific volume with each SV. For this purpose, we generated the Voronoi tessellations for each SV cluster (Fig. 4a); here SVs are colored in green and the surrounding red structure is the Voronoi Tessellation (see methods). In Fig. 4b, we represent some individual Voronoi volumes with their respective SVs in green. For each SV we measured the SV-associated volume; this is the remaining volume after subtracting the SV volume from the Voronoi volume (Fig. 4c). The distribution of the SV-associated volumes is presented in Fig. 4d. Under LTP conditions, SVs have associated more volume, in accordance with the loss of SVs and conservation of the SV cluster volume.

**Figure 4.**
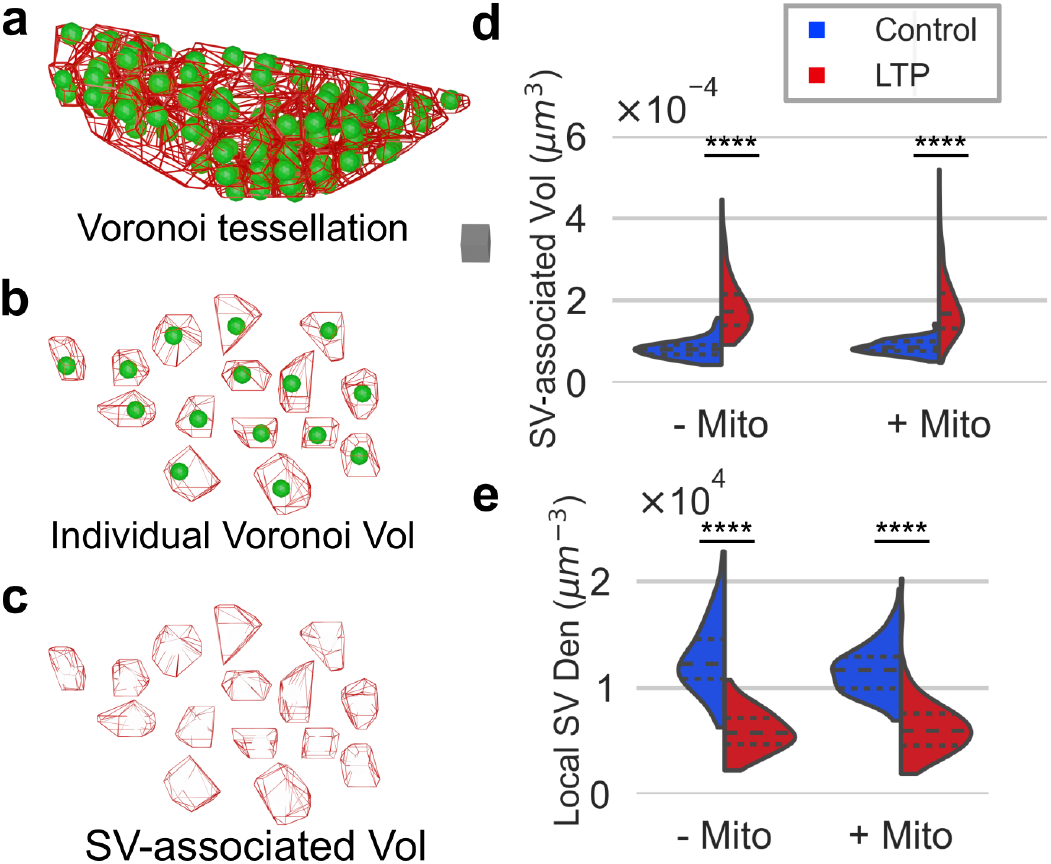
The SV-associated volume for each SV increases after LTP induction. (a) Voronoi tessellation for one representative SV cluster in red and SVs in green. Scale cube = 0.001 µm^3^. (b) Individual Voronoi volumes for some SVs in the cluster. (c) The SV-associated volume is the remainder of subtracting the SV vol from each Voronoi volume (see methods). (d) The SV-associated volume increases after LTP induction. (e) The local SV density (Local SV Den) reduces after LTP (i.e. 1 divided by the SV-associated volume) for boutons without and with mitochondria under control and under LTP conditions. Violin plots show quartiles 25th, 50th, and 75th. Significant differences are indicated as ^*^p < 0.05, ^**^p < 0.01, ^***^p < 0.001, and ^****^p < 0.0001. Medians, standard deviations, and p-values are in Table 1.

This partition also allows us to quantify the local SV density, i.e. one divided by the SV-associated volume (Fig. 4e). After LTP induction, the local SV density is reduced in relation to control conditions (p < 8.00 ×10^−20^ -Mito and p < 6.00 ×10^−24^ +Mito).

As we will show in the next section, the distributions of Voronoi volumes contain information about the spatial organization of SVs; as well as the relationship between the global and local SV densities.

### Spatial organization of SVs at synapses

The Voronoi tessellations contain information about the spatial organization of the set of points in space. Fig. 5a shows a representative example of the 2D Voronoi tessellation for a set of randomly, clustered, and uniformly distributed points. The distribution of Voronoi sizes (areas in 2D and volumes in 3D) differs depending on the spatial organization of the points. Smaller Voronoi areas are observed when points are clustered in space (Fig. 5a). The distribution of Voronoi volumes has been employed in the past to identify clusters of proteins in microscopy images [19, 20]. It provides a tool for transforming questions about the arrangements of points into ones about areas and volumes [21].

**Figure 5.**
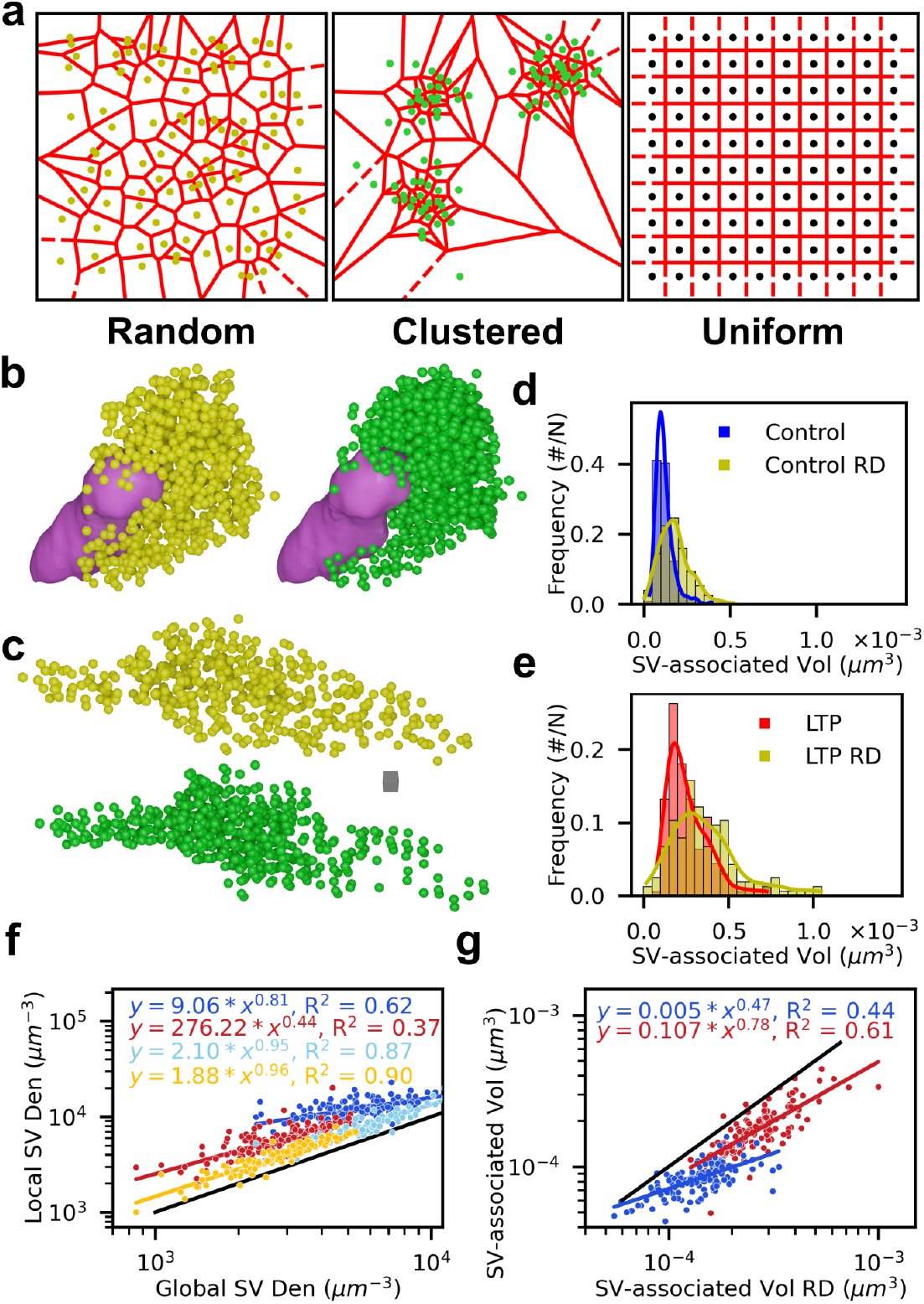
Spatial organization of SVs at synapses. (a) 2D Voronoi tessellation for a set of points randomly distributed, clustered, and uniformly distributed. Points are in yellow, green, and black, respectively, and tessellations are in red. The distribution of Voronoi sizes differs depending on the spatial organization of the points. Smaller Voronoi sizes are observed when points are clustered. (b) Representative examples of a SV cluster under control conditions in green and randomly distributed SVs in yellow. Scale cube = 0.001 µm^3^. (c) Same as in (b) for a SV cluster under LTP conditions in green and randomly distributed SVs in yellow. (d-e) For each bouton represented in (b-c), we plot the distributions of SV-associated volumes. Larger volumes are observed for randomly distributed SVs. (f) Local vs. global SV densities at each bouton under control in blue and under LTP conditions in red, to compare the values to a uniform distribution we plot a line with slope and intercept equal to one in black. The local SV density is larger than the global density, implying that the SV-associated volumes are smaller under control and LTP conditions than for a uniform distribution. For the randomly distributed (RD) SVs the values are more similar to the uniform distribution but always below the values for the original SVs, under control conditions in light blue and under LTP conditions in yellow; which implies the original distribution of SVs is not random, and is more compact under control conditions. (g) SV-associated volume for the original vs. randomly distributed SVs. After the induction of LTP, the distribution of SV is more similar to a random distribution.

We compare the spatial distribution of the original SVs with randomly distributed (RD) SVs at each bouton, keeping the volume of the SV cluster constant (see methods). Figs. 5b-c show for two representative boutons how the distributions of SVs in space change from the original distributions (in green) to the randomly distributed (in yellow), under control (Fig. 5b) and under LTP conditions (Fig. 5c). Under control conditions (Fig. 5d) the distribution of SV-associated volumes is concentrated towards smaller values for the original SVs (in blue), which implies SVs are more clustered than in the randomly distributed example (p-val < 7 × 10^−33^). Similarly, under LTP conditions the SV-associated volumes of the original SVs are smaller than randomly distributed SVs (Fig. 5e, p-val < 2 × 10^−8^).

If we think in terms of densities instead of volumes. The global and local densities are equal for a uniform distribution of points. While they differ for a randomly or clustered distributed set of points. In Fig. 5f we compare the local SV density to the global SV density under control (in blue) and under LTP conditions (in red); to compare the values to a uniform distribution we plot a line with slope and intercept equal to one in black. The local SV density is larger than the global SV density, implying that the SV-associated volumes are smaller under control and LTP conditions than for a uniform distribution, and in particular under control conditions, the local density is the largest. This suggests that SVs are tightly packed under control conditions. We also show the local and global SV densities for randomly distributed (RD) SVs under control in light blue and under LTP conditions in yellow (Fig. 5f). The local density is larger than the global density also for SV randomly distributed but the difference is smaller than for the original distributions.

To compare the SV-associated volume for each bouton with the SV-associated volume of the randomly distributed SVs, in Fig. 5g we plot the median of each distribution: in the y-axis the medians for the original distributions and on the x-axis the medians for the randomly distributed SVs. As before we added a line with slope and intercept one to compare the medians. The medians of the randomly distributed SVs are always larger than for the original distributions of SVs. One conclusion we can derive from this analysis is that the distribution of SVs is not random. Moreover, under LTP the medians are more similar to the values of the randomly distributed SVs (closer to the black line). Therefore, under LTP, the distributions are more similar to the randomly distributed SVs.

### Potentiated synapses have lower SV density, larger distance between SVs, and are more dispersed from the SV cluster’s center

All the previous analyses showed us the SV clusters present different structural features after the induction of LTP. Here, we assessed how these quantities vary together and developed methods to identify potentiated synapses based on structural features. For this, we use principal component analysis (PCA) with several of the features measured for the SV cluster –presented in the previous sections. We used the distances to neighboring SVs, the global SV density, the local SV density, the dispersion and the bouton surface area to separate the boutons into two clusters.

Fig. 6a shows how the distances to neighboring SVs increase as the global SV density decreases under LTP. Boutons with larger distances to neighboring SVs are those that have been potentiated. To further test this assumption, we analyzed reconstructions of axons treated with APV to block LTP. Fig. 6b shows the distances to neighboring SVs versus the global SV density for axons treated with APV stimulated under control and under LTP conditions. As expected, these results mainly overlap with the measured quantities under control conditions.

**Figure 6.**
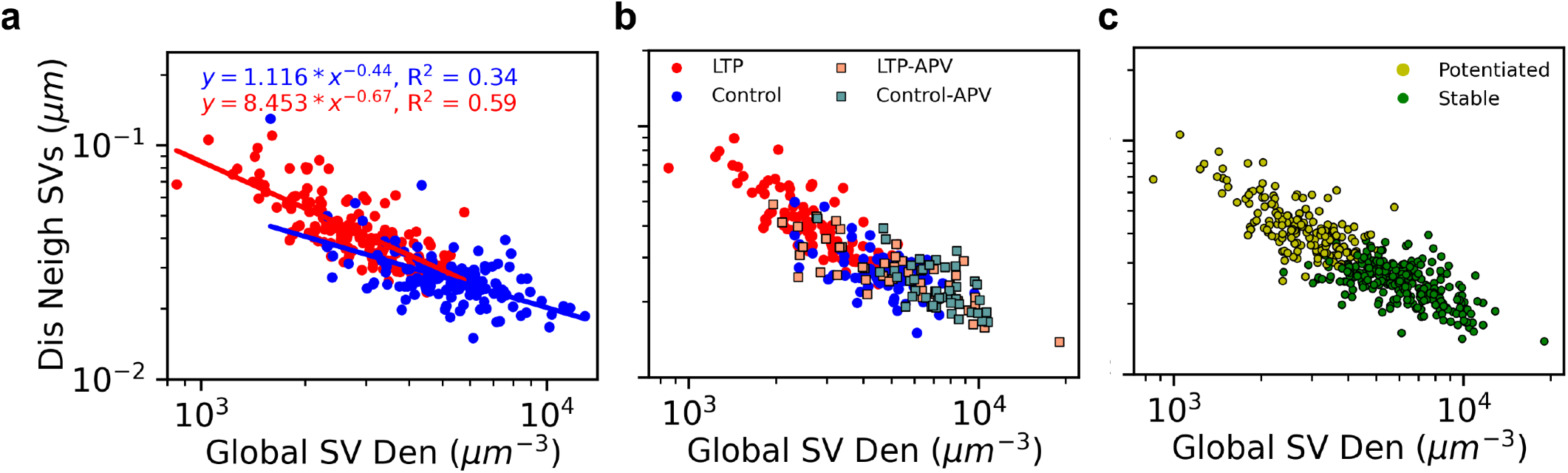
Potentiated synapses have lower local and global SV densities, larger distances to neighboring SVs, and are more dispersed from the cluster’s center. (a) Distances to neighboring SVs vs global SV density under control conditions in blue and under LTP conditions in red (b) Same as in (a) but we included measurements from reconstructions from brain slices treated with APV, where LTP has been blocked. (c) Two populations of boutons are identified using five features of the SV cluster employing the K-means clustering algorithm, using the first two principal components of PCA. The potentiated population has a lower SV density, higher distance to neighboring SVs, higher SV-associated volume, and SV are more dispersed from the center while the opposite occurs in the other population.

To cluster boutons with similar properties, we first use PCA to identify the variables that better capture the variability of the data. All the data were scaled before the analysis, centered to zero, and normalized to have a standard deviation of one. We also log transform the data before doing PCA. The first two principal components account for 90 % of the variance. Therefore we use these two components as input for the clustering method. We cluster the boutons in two groups: potentiated and stable boutons (Fig. 6c). We employed the K-means algorithm to cluster the boutons. We mapped the labels obtained after kmeans to the original values, of the distances to the neighboring SVs and the global SV density (Fig. 6c).

### The Local SV density is a volume-independent property

Next, We ask how the SV cluster volume varies with the bouton volume. We found the SV cluster volume scales with the bouton volume in all conditions in a similar manner (Fig. 7a).

**Figure 7.**
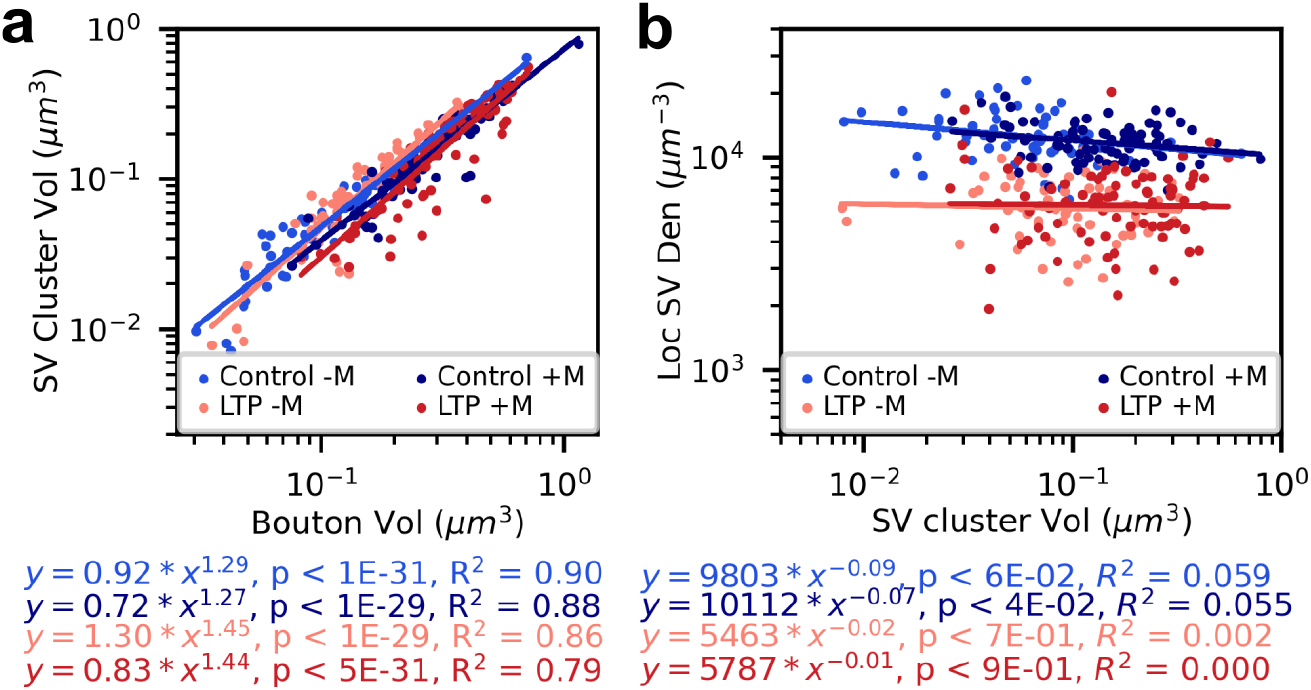
The local SV density is a volume-independent property. (a) The SV cluster scales with the bouton volume; similar scaling properties are observed for boutons with (+M) and without mitochondria (-M). Under control conditions in light blue (-M) and dark blue (+M), and under LTP in salmon (-M) and red (+M). (b) On the contrary the local SV density is an intrinsic property of the system, that remains constant for all SV cluster volume under a given condition, but it changes after the induction of LTP. The local SV density remains constant over three orders of magnitude of the SV cluster volume, which suggests it might be under tight regulation. This indicates the volume associated to each SV is tightly regulated and changes after the induction of LTP. Values of the slope, intercept, p-value, and R^2^ are in the text below.

We evaluated how the local SV density relates to the SV cluster volume (Fig. 7b). We found the local SV density remains invariant under control or LTP conditions over a wide range of the SV cluster volumes. This is observed in boutons with and without mitochondria, suggesting the SV density is likely subject to strict regulatory mechanisms. The local SV density remains also invariant to the bouton volume.

Next with a simple computational model and theoretical calculations, we explored the potential implications of the change in the SV density from control to LTP conditions. The model considers the diffusion of SV in 3D space and assumes different diffusion coefficients (or viscosities) for the SVs in the axon and in the SV cluster. Due to the different diffusion coefficients a gradient in the density (or concentration) can be maintained. A concentration gradient can be held by a difference in the diffusive properties of the medium. Assuming this is the case between the axon and the SV cluster, then knowing the diffusion coefficient in one of the mediums we can estimate the diffusion coefficient in the other medium. We calculated the diffusion coefficients with a simple theoretical calculation. At equilibrium, the number of times SVs hit (N_hits_) at the surface of the SV cluster from inside and outside should be the same. The number of hits has been calculated [22] for a given concentration (C), diffusion coefficient (D), Avogadro number (N_A_), time interval (Δ*t*) and surface area (Area) of the SV cluster:

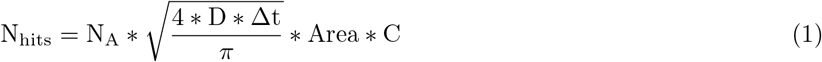

Therefore, we can compare first the number of hits from inside the SV cluster to the number of hits from the axon side under control conditions. The diffusion coefficient of SVs in the cluster has been measured (D_Contol_= 0.05 µm^2^ s ^−1^ [13]. In this manner, we can estimate the diffusion coefficient of the SVs in the axon (Table 2). This also allows us to estimate the change in the diffusion coefficient from the axon to the SV cluster.

**Table 2:** Parameter values employed in the computational model.

| Property | Axon | SV Cluster |
| --- | --- | --- |
| Volume ($\mu\text{m}^3$) | 0.583 | 0.128 |
| Density of SVs ($\mu\text{m}^{-3}$) | 220 | Control: 4688 & LTP: 2344 |
| Diffusion Coefficient ($\text{cm}^2 \text{s}^{-1}$) | $2.244 \times 10^{-5}$ | Control: $5 \times 10^{-10}$ & LTP: $2 \times 10^{-9}$ |

We can relate the diffusion coefficient with the viscosity of the SVs, which is three orders of magnitudes higher than in the axon, similar to what has been measured in other condensates [23]. Second, assuming the conditions at the axon remain stable after LTP induction, we can thus compare the number of hits inside the SV cluster for control and LTP conditions. Since the reduction in SV density is approximately half after LTP induction, this implies the ratio of the diffusion coefficients has to be one quarter (equation (1)); which means the diffusion coefficient after LTP induction is four times the one under control condition, which summarizes to D_LTP_ = 0.2 µm^2^ s^−1^. To recapitulate, with our simple theoretical calculation that assumes different diffusion coefficients in the axon and in the SV cluster we can predict an increase in the diffusion coefficient of the SV cluster after LTP induction.

We tested the prediction using a 3D reaction-diffusion model of the axon and the SV cluster, with SVs diffusing in 3D space (Fig. 8b). The model was implemented in MCell4, an agent-based reaction-diffusion simulator [24]. Fig. 8b shows how the number of SVs is maintained due to the difference in diffusion coefficients between the axon and the SV cluster (blue and black are single traces). At 2.5 seconds we change the diffusion coefficient of the SVs in the cluster, and after some seconds the system goes back to equilibrium with now half of the number of SVs in the cluster. We show that a change in the viscosity of the media can account for the measured change in the density of SVs in the cluster.

**Figure 8.**
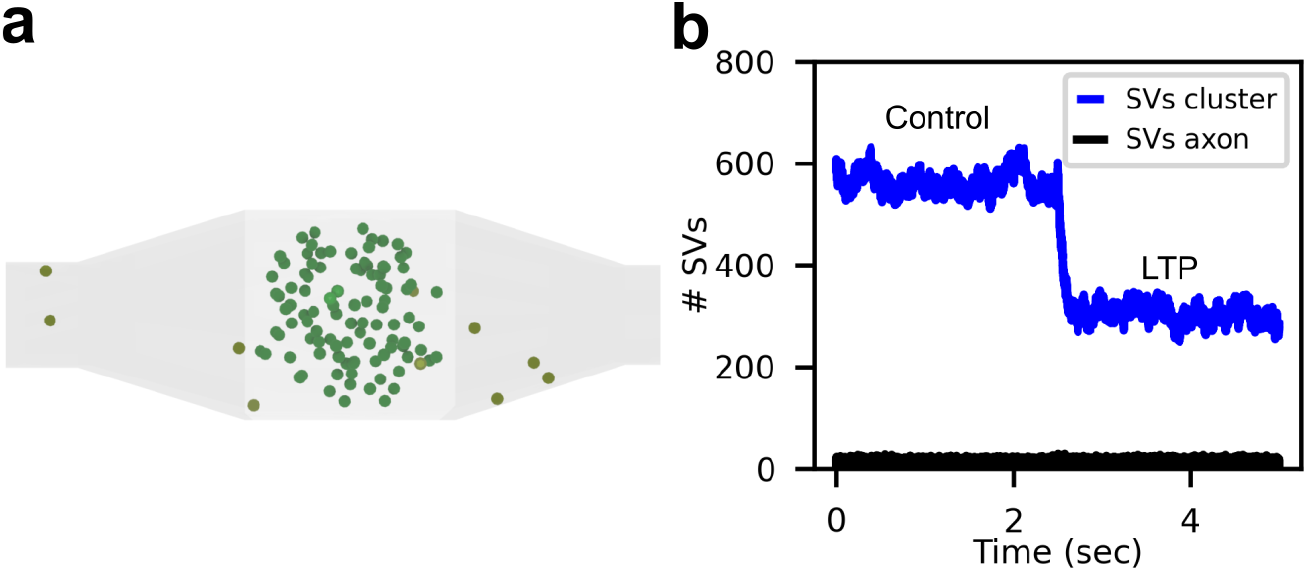
A simple computational model assuming different diffusion coefficients in the axon and the SV cluster was used to predict the implications of the change in the SV density after LTP induction. (a) In the computer simulations we assume SVs diffuse with different coefficients in the axon in yellow and in the SV cluster in green. In this manner, a gradient in the density is maintained. In the computer simulations, we assume the diffusion coefficient in the SV cluster changes after the induction of LTP. This translates into a change in density in the SV cluster. (b) Before 2.5 seconds, the system is at equilibrium with SVs diffusing in the SV cluster and the axon with values similar to those in control conditions; at 2.5 seconds, we change the diffusion coefficient in the SV cluster, and the system relaxes to the new equilibrium, with lower SV density.

## Discussion

One of the most challenging and novel problems of our times in synaptic physiology is the formation of biological condensates [10, 25]. Accumulating evidence shows this is a ubiquitous process in biology [26], and particularly in synaptic physiology [10]. Dysregulation of condensate formation in neurons has been associated with several pathological conditions [23, 27]. Therefore, understanding this phenomenon has implications for health and disease. Condensates of synapsin I have been implicated in clustering lipid vesicles in vitro [12] and SVs in vivo [13, 14]. For instance, it was shown that the disorder domain of the phosphoprotein synapsin I is needed for establishing the formation of SVs condensates [14, 12]. Several phosphoproteins participate in regulating the SV cluster [15, 16, 17]. In particular, the phosphorylation of synapsin I by protein kinases after synaptic activation dissociates synapsin from SVs and seems to control the mobilization of SVs [15, 28]. In this context, we asked two questions: Is the SV cluster altered after LTP induction in CA1 hippocampal boutons? And if it is, what are the functional implications of these changes?

Our analysis reveals that the structure of the SV cluster is altered after 2h of TBS. We found the distances to neighboring SVs increases, SVs are more dispersed from the cluster’s center, the cluster-associated volume is maintained after LTP induction, and the SV-associated volume increases, while other properties remain unaltered as the position of the SV cluster center and the distance from the SVs to the AZ. Remarkably, some of these properties are independent of the presence of mitochondria at the bouton, such as the distance to neighboring SVs or the SV-associated volume. While other properties differ in boutons with mitochondria, such as the dispersion of the SVs from the cluster’s center or the cluster-associated volume.

The SV cluster has been speculated to act as a key regulator and organizer of the synaptic composition, dynamics, and homeostasis [29]. It has been suggested that the SV cluster is regulated to control the protein copy numbers at synapses [29]. In this work, we revealed that the local density of SVs is a volume-independent property, i.e., all synapses, big or small, have the same SV density under the same conditions. The local SV density is kept constant over three orders of magnitudes of the SV cluster volume for a given condition (Fig. 7b), which suggests this intensive property of the SV cluster must be under strong regulation.

Biological condensates are thought to undergo phase transitions, in analogy to how matter changes between different states [30]. A phase transition is the physical change process between two system states. For instance, the most common example is the transition of water from a liquid to a solid state. In this transition, the temperature acts as the control parameter governing the change from liquid to solid or vice versa. During such transitions, intensive properties of the system change. Post-translational modifications are an essential control mechanism of phase transitions [26]. For instance, the degree of phosphorylation of Nephrin governs the transition of one of such condensates through the interaction with protein ligands called Nck and N-WASP and helps filaments of actin to form [30]. For the SV cluster, the concentrations of synapsin I and synuclein have been suggested to act as a control parameter of such a transition [31] affecting in this manner the phase of the condensate, density, and volume of the SV cluster. Here, we show that an intensive property of the SV cluster changes after LTP induction which suggests an analogy with phase transitions. We provide further evidence of such a transition, showing that 2 hours after TBS stimulation, the SV cluster is more dispersed and less dense than under control conditions. Furthermore, we have recently shown that SVs permanently fuse during LTP to expand the bouton surface area [6]. Together with the results presented in this paper, we show that the SV cluster is undergoing a transition from a compact to a loose organization, which has functional implications. During this transition, the recycling of SVs is inhibited. Further investigation would allow for a deeper understanding of the functional consequences of such a transition.

In a recent publication [6], we showed that the bouton surface area expands after 2 hours of LTP induction. To increase the synaptic strength or efficacy, not only lipids are needed but also proteins. Therefore, we can speculate that perhaps after LTP induction, SVs have more associated volume to bind or allocate more proteins needed to expand the synaptic efficacy of the synapse. It could also be noted that many condensate properties could affect reactions within them. For instance, molecular crowding, the reduction in available volume due to high molecular concentration, can influence the regulation and binding affinity and, in turn, alter enzymatic activities [26].

The size of subcellular organelles is tightly controlled in single cells [32, 33] and is dysregulated in diseases such as cancer. Scaling relations are important because they suggest possible size control mechanisms. Scaling relations can arise from different mechanisms, either genetically programmed regulatory pathways or relative growth or physical constraints [33]. In mammalian cells, the cytoplasm fraction occupied by mitochondria is relatively constant [34]. In the brain, scaling properties of subcellular organelles with the bouton volume were extensively studied for different cell types [35, 36, 37]. Here, we provide evidence that the SV cluster volume is also under tight regulation; it scales with the bouton volume in a similar manner under control and LTP conditions (Fig. 7a) as well as other properties of the SV cluster such as the number of SVs or the total SV volume which also scale with the bouton volume. Mitochondrial volume on the other hand more loosely correlates with the bouton volume (Fig. 1k); and a slight reduction in the correlation is observed after LTP induction, due to small boutons having more mitochondria (Fig 1k).

We acknowledge that our approach has some limitations. First, the section thickness of the EM images is approximately 60 nm, larger than the diameter of the SVs (40 nm). To test how our results depend on the section thickness, we repeated our measurements randomly, shuffling the center of the SVs by ± 31 nm in the z-direction. The percentage error is smaller than 3%, and the differences between control and LTP are maintained (p-val < 1.2 ×10 ^−4^ for boutons without mitochondria and p < 9 ×10 ^−7^ for boutons with mitochondria). In the future, we plan to do a more comprehensive study using newly developed high-resolution large-field electron tomography to reconstruct SVs in 3D more accurately.

We predict the implications of a change in SV density at synapses with a simple computational model and theoretical calculations. The model, though phenomenological, has allowed us to estimate the change in diffusion coefficient (or viscosity) from the axon to the SV cluster, a finding that is in order of magnitudes similar to what was measured in other condensates [23]. This approach led to the prediction of an increment in the mobility of SVs in the cluster after LTP induction. When we compared the distributions of SV-associated volumes between the originals and randomly distributed SVs, we observed that the SV-associated volumes after LTP induction were more similar to the randomly distributed SVs. If SVs are gaining mobility, it is expected that the distributions of SV-associated volume will be more similar to the randomly distributed SVs.

This work employs multiple approaches to characterize the clustering of SVs and reveals the modifications to the SV cluster after LTP induction. In the future, it would be fundamental to determine whether a similar response is observed in young and adult animals. Do mature and young animals respond in the same manner? It would be critical to characterize these transitions during the natural aging process. Lastly, developing a multiscale computational model that captures the interaction of synapsin I and synuclein with SVs will be an invaluable tool to explore the functional implications of condensate formation on synaptic transmission. Moreover, one promising aspect of the modeling approach is to test which are the fundamental protein-protein interactions that drive the condensate formation. This work will be the foundation for future models of the SVs cluster in synapses. Understanding the mechanism by which SVs cluster is fundamental to synaptic transmission, learning, and memory formation.

## Methods

### Slice preparation, electrophysiology, and microwave fixation

Brain slices from the middle of the rat hippocampus were prepared as previously described [7]. Two concentric bipolar electrodes were lowered into the middle of stratum radiatum in area CA1 [7] separated by 500 µm, stimulating independent axons. Control stimulation (one pulse every two minutes for 40 minutes) was delivered to one of the electrodes and TBS to the other one (8 trains of 10 bursts at 5 Hz of four pulses at 100 Hz delivered 30 sec apart). 2 hours following TBS, the slices were fixed, processed, and imaged.

### 3DEM Reconstructions

The EM images were visualized in the new version of Reconstruct [38] (PyReconstruct) [39]. 3D reconstructions of axons and mitochondria were generated using the NeuropilTools addon for Blender (version 2.79) [40]. The program Contour Tiler [41], which is integrated into NeuropilTools, was employed to create triangulated surface meshes from the 2D traced contours. The positions of the center of the SVs was assessed manually.

### Total Vesicle Volume

A subset of the SVs was traced from each dataset to estimate the total SV volume. The circumference of a subset of SVs was measured, and the radii were calculated, assuming circular SVs. Afterward, bootstrapping analysis was used: 10.000 bootstrap samples were selected to estimate the total SV volume and its error for each bouton.

### Morphometric Analysis

Several Python scripts were developed to analyze the distribution of SVs in 3D in Blender. The scripts utilized Python 3.8, Scipy version 1.5.2, and Numpy version 1.19.2. All the measured quantities are stored in a csv file in the repository, along with the generated scripts (in the folder named scripts_blender). To further analyze and plot these measurements another set of scripts was developed, using Pandas version 1.1.5, Matplotlib version 3.3.2, Seaborn version 0.11.1, and Sklearn version 0.23.2. All the associated material can be found in the repository (in the folder scripts_figures).

Statistical significance between distributions was assessed using the nonparametric Mann–Whitney–Wilcoxon test implemented in SciPy.

### Distance to neighboring SVs

To compute the distance to neighboring SVs, we generated the Delaunay triangulation for each SV cluster. For a given set of points, the Delaunay triangulation is such that no point in the set is inside the circumcircle of any triangle in the triangulation. Neighboring SVs are connected vertices in the Delaunay triangulation. The Euclidean distance between neighboring SVs was calculated and the mean distance was computed for each SV. The triangulation was generated using the associated routine in Scipy, through a Python script written for Blender 2.79. Finally, the median was computed from these distributions of distances, which is the final quantity reported for each bouton. We subtracted two times the median radius of the SV to compute the final distance, which we refer to as the distance to neighboring SVs throughout the manuscript.

### Convex Hull

For a set of points, the convex hull or envelope is the smallest convex set that contains it. The convex hull for the SV cluster was computed using the routine ConvexHull in Scipy. Afterward, the associated 3D mesh was generated in Blender. The intersection between mitochondria and the axon with the convex hull was computed using the Blender “Boolean Difference Modifier”. The intersection was subtracted from the convex hull volume. Finally, the convex hull volumes were calculated employing the meshalyzer plug-in in CellBlender to calculate surface areas and volumes [40].

### Voronoi tessellation

The volumes associated with each SVs were determined using the Voronoi tessellation for the SV cluster. A routine in Scipy was used for this purpose, and the 3D volumes were generated in Blender. The intersection of each volume with the cluster’s Convex Hull was assessed. The plug-in meshalyzer in Cellblender was used to compute the Voronoi volumes. Only the unaltered Voronoi volumes after calculating the intersection with the Convex Hull were used. The final value reported in the manuscript is the Voronoi volume minus the SV volume, referred to as the SV-associated volume.

The distribution of Voronoi areas (in 2D) and volumes (in 3D) provide information about the spatial distribution of the components in space [21]. When points are regularly distributed, the Voronoi volumes do not vary much. But when points are clustered, smaller volumes are present. For instance, the distribution of Voronoi areas (in 2D) has been used to identify clusters of proteins from super-resolution images [42, 43].

### Center of the SV cluster and dispersion

The center of the SV cluster was calculated with a Python script in Blender, and the Euclidean distances from the SVs to the center of the SV cluster were calculated. The median radius of the SV was subtracted to compute the final distance.

The standard deviation (σ_xyz_) of the SVs to the cluster’s center was assessed. This quantity measures the dispersion of the SVs from the SV cluster center (equation (2)), here 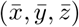 are the coordinates of the SV cluster center, (*x*_*i*_, *y*_*i*_, *z*_*i*_) are the coordinates of each SV and N is the total number of SVs in the cluster. In the literature, it has been referred to as the standard distance.

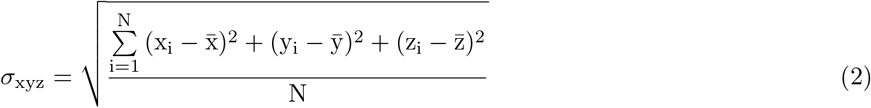

### Random Distribution of SVs in the cluster

To compare the spatial distribution of SVs in space, we generated a new distribution for each cluster following a Poisson distribution. For this, we used the 3D-reaction diffusion simulator MCell4 [24]. For each SV cluster, the same number of SVs in the bouton were distributed inside the Convex Hull. MCell4 gave the new position of each SV and subsequently, we imported it in Blender to generate the new position of the SV in 3D. The same methods described before were applied to the randomly distributed SVs and the SV-associated volume for each SV was assessed.

### A computational model with different SV densities and diffusion coefficients

A diffusion model of the SVs was built in MCell4, assuming distinct densities of SVs and diffusion coefficients in the axon and the SV cluster. Inside the SV cluster, the density of SVs and the diffusion coefficient were considered high, while outside were considered low (see Table 2 for parameter values). The change in the SV density was used to estimate the change in the diffusion coefficient of SVs in the cluster after LTP induction (parameters also shown in Table 2).

## Author contributions statement

G.C.G., T.M.B., L.K., K.M.H. and T.J.S. conceived and designed the research; G.C.G., T.M.B. developed software, G.C.G. L.K., and P.B. analyzed the data, G.C.G., T.M.B., L.K., K.M.H., and T.J.S. wrote the paper. All authors agree on the contents of the manuscript.

## Conflict of interest statement

All authors declare no known or potential conflict of interest including any financial, personal, or other relationships with other people or organizations that could inappropriately influence, or be perceived to influence, their work.

## Data Availability

All the software generated during the current study is available in the GitHub repository: https://github.com/guadagar/axon_cyto_git.

## Acknowledgements

We thank Prof. Mary B. Kennedy for insightful discussions on the formation of condensates in synapses. This work was supported by NSF Technology Hub Award No. 1707356 and NSF NeuroNex2 Award No. 2014862

## Notes

### Competing Interest Statement

The authors have declared no competing interest.

