## Supplementary material for "The presynaptic vesicle cluster transitions from a compact to loose organization during long-term potentiation"

The median distances from the SVs to the AZ are similar for single synaptic boutons (SSB) and multisyanptic boutons (MSB) (Fig. 1) under control and under LTP conditions.

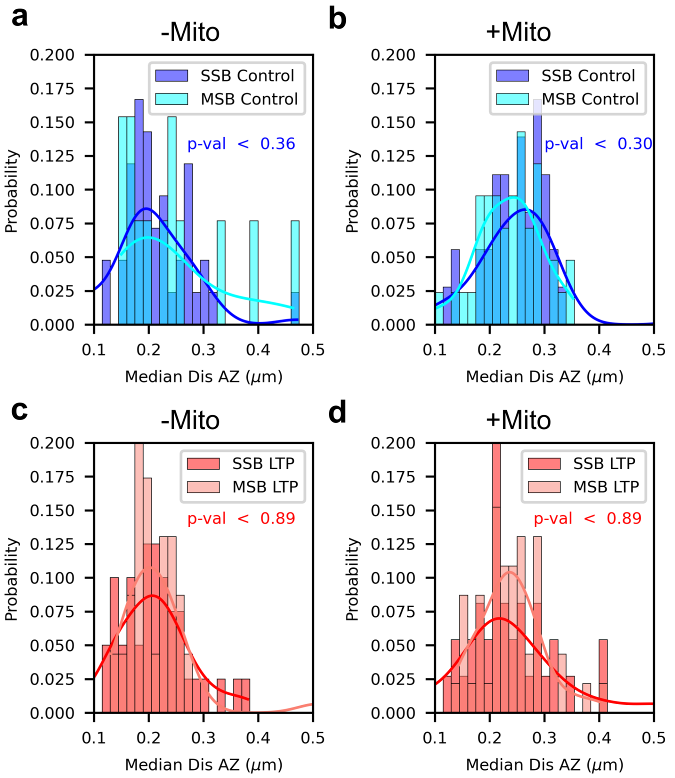


Figure 1: The median distances to the active zone (AZ) are similar in single synaptic boutons (SSB) and multisynaptic boutons (MSB) (a) under control conditions without mitochondria and (b) with and under LTP conditions (c) without mitochondria and (d) with mitochondria.

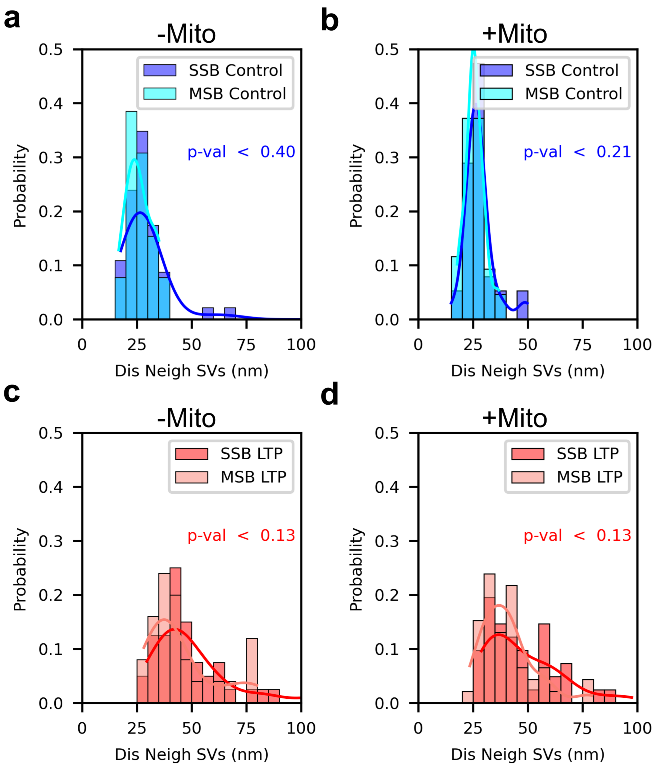


Figure 2: The mean distances to neighboring SVs are similar in single synaptic boutons (SSB) and multisynaptic boutons (MSB) (a) under control conditions without mitochondria and (b) with and under LTP conditions (c) without mitochondria and (d) with mitochondria.
